# Differential BUM-HMM: a robust statistical modelling approach for detecting RNA flexibility changes in high-throughput structure probing data

**DOI:** 10.1101/2020.07.30.229179

**Authors:** Paolo Marangio, Ka Ying Toby Law, Guido Sanguinetti, Sander Granneman

## Abstract

Combining RNA structure probing with high-throughput sequencing technologies has greatly enhanced our ability to analyze RNA structure at transcriptome scale. However, the high level of noise and variability encountered in these data call for the development of computational tools that robustly extract RNA structural information. Here we present diffBUM-HMM, a noise-aware model that enables accurate detection of RNA flexibility and conformational changes from high-throughput RNA structure-probing data. DiffBUM-HMM is compatible with a wide variety of high-throughput RNA structure probing data, taking into consideration biological variation, sequence coverage and sequence representation biases. We demonstrate that, compared to the existing approaches, diffBUM-HMM displays higher sensitivity while calling virtually no false positives. DiffBUM-HMM analysis of *ex vivo* and *in vivo* Xist SHAPE-MaP data detected many more RNA structural differences, involving mostly single-stranded nucleotides located at or near protein-binding sites. Collectively, our analyses demonstrate the value of diffBUM-HMM for quantitatively detecting RNA structural changes and reinforce the notion that RNA structure probing is a very powerful tool for identifying protein-binding sites.

## Introduction

Understanding the structure of an RNA is key to unravel its *in vivo* function, and it is also highly relevant to biomedicine, drug discovery and synthetic biology (1–4). Recent years have witnessed a blossoming of high-throughput methods that couple next-generation sequencing with biochemical assays to “probe” the structure of thousands of RNA molecules simultaneously, including whole transcriptomes. (5–16). The majority of these biochemical assays use reagents such as SHAPE (Selective 2‘-hydroxyl acylation analyzed by primer extension) reagents (5–7, 16–20) and dimethyl sulfate (DMS) (8, 10, 21). These chemicals modify the 2‘-hydroxyl (OH) group of riboses or bases of flexible/single-stranded nucleotides, respectively, and the sites of modification can be detected by performing a reverse transcription (RT) reaction. A major advantage of using chemical probes is that some are also highly effective for probing RNA structure in living cells (16, 18, 22–24), making it possible to compare *in vivo* and *in vitro* structures, and reveal potential protein-binding sites (16, 22). Depending on the RT enzyme and the reaction chemistry used, the modification either causes the RT enzyme to terminate transcription, resulting in truncated cDNAs, or to skip the adduct, frequently introducing mutations (SHAPE-MaP; (5, 21)). Following on from this, the site and degree of nucleotide modification can be extracted from NGS data by quantifying how frequently the RT terminated at a given nucleotide position (6, 8–10) or by calculating mutation frequencies for each nucleotide (5, 21). Although NGS has a number of unprecedented advantages in terms of sensitivity and the number of molecules that can be analyzed simultaneously, the analysis of the resulting data is not trivial and exhibits significant challenges. Depending on the cDNA library preparation method used, biases in sequence representation and read coverage can be introduced (25), and there can also be quite significant inter-replicate variability in untreated (control) and treated samples (26). To specifically address these issues, we recently developed a probabilistic modelling pipeline called beta-uniform mixture hidden Markov model (BUM-HMM) (27). One of the strengths of BUM-HMM is that it analyzes the inter-replicate variability of samples in the treatment and control pools. Moreover, it adopts an empirical statistical analysis method that obviates the need of conventional data correction and normalization techniques that are used in the majority of the analysis pipelines. Although BUM-HMM generates statistically sound estimates of nucleotide accessibility at the nucleotide level, its probabilistic output does not represent an absolute value that quantifies the degree of accessibility of RNA at a particular nucleotide. Therefore, it is not immediately usable for differential analyses between different treatments.

The ability to accurately detect nucleotide regions that under diverse conditions differentially react with RNA structure probing reagents is of great importance to researchers. As a consequence, the last few years have seen an increase in the development of a number of bioinformatics tools to detect differentially reactive nucleotides (DRNs) in RNA structure probing datasets. Among the available tools are classSNitch (28), PARCEL (29), RASA (30), deltaSHAPE (22), StrucDiff (31), and the recently published dStruct (32). In particular, dStruct has been shown to perform best by recording the lowest false positive rate, while offering compatibility with a wide range of existing RNA structure probing datasets. However, one possible limitation of dStruct is that the pipeline uses a variety of statistical tests to predict DRNs. As a result, dStruct corrects for multiple hypothesis testing, which likely makes it conservative with its predictions. Hence, we reasoned that a method that does not rely on statistical tests but rather on a model and posterior probability, such as BUM-HMM, would be preferable, because it would be inherently less vulnerable to problems associated with multiple hypothesis testing. In addition, dStruct imposes normalization and outlier elimination strategies on quantitative data to generate a reactivity profile for each nucleotide. Since the distribution of quantitative data often differs between probing experiments, such procedures might result in useful data being removed. In contrast, the BUM-HMM model uses only the raw counts for each nucleotide (i.e. read coverage and either total RT drop-offs or mutation counts). It also employs empirical statistical analyses that preserves the independent distribution of each dataset whilst being robust to outliers. To test whether the BUM-HMM algorithm could be useful for detecting DRNs, we extended the model to develop diffBUM-HMM (differential BUM-HMM). We used diffBUM-HMM to compare a number of publicly available RNA structure probing datasets and benchmarked the tool against dStruct (32). Similar to dStruct, diffBUM-HMM effectively identified DRNs in the datasets, however, consistent with our hypothesis, it exhibited higher sensitivity and, like dStruct, has a very low false positive rate. Because diffBUM-HMM is compatible with a wide variety of high-throughput RNA structure probing methods, it should be of general interest to the RNA community.

## Results

### DiffBUM-HMM Model

DiffBUM-HMM is a natural extension of BUM-HMM (Fig. 1). An intermediate step of BUM-HMM is the computation of an empirical *P* value for each treatment-control comparison at each nucleotide position. Each empirical *P* value is then passed onto a hidden Markov model. BUM-HMM has a *hidden state h_t_* (*t* = 1, 2, 3, …, *T* for *T* nucleotides) representing the true binary state of the *t*-th nucleotide (M - modified by the probe; or U - unmodified by the probe), and the observed variable *v_t_*, which is the empirical *P* value at that position. For diffBUM-HMM, the *hidden state* is expanded to take on four potential values instead of two: nucleotide is unmodified in both conditions (UU; *hidden state* 1); nucleotide is unmodified in the 1^st^ condition but modified in the 2^nd^ (UM; *hidden state* 2); nucleotide is modified in the 1st condition, but unmodified in the 2^nd^ (MU; *hidden state* 3); nucleotide is modified in both conditions (MM; *hidden state* 4). In turn, the observed variable *v_t_* at each state is now represented by two *P* values rather than one. As the *hidden state* can take on four possible values, extending BUM-HMM to diffBUM-HMM entails increasing the size of the transition matrix from 2×2 to 4×4 and adapting its values. While in principle an EM algorithm could be used to identify directly this transition matrix from data, we found that adapting the original BUM-HMM heuristic values to diffBUM-HMM by assuming independence of the two conditions yielded good results. A sensitivity analysis confirmed the validity of this approach (Supplementary Fig. S1).

**Fig. 1.**
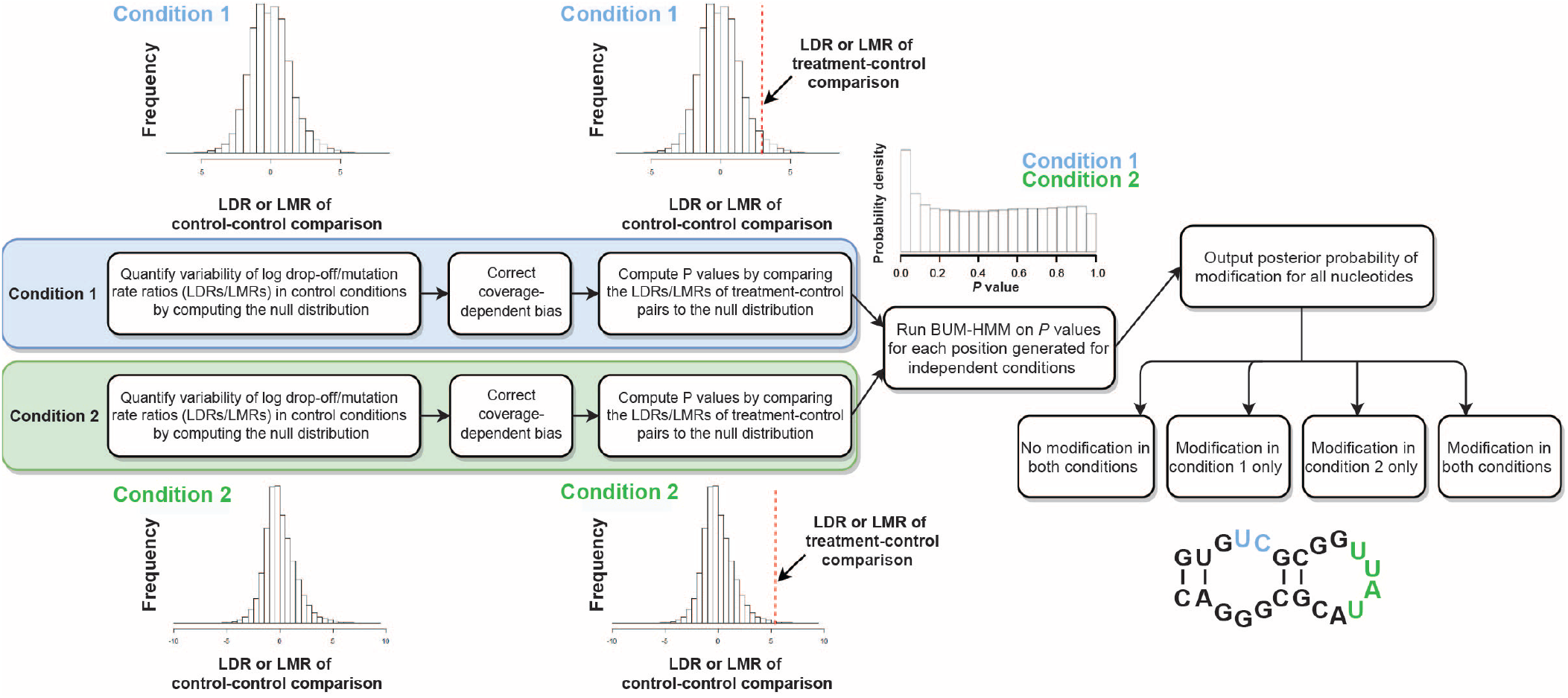
Overview of the diffBUM-HMM computational analysis pipeline. For each experimental condition (e.g. Condition 1 and 2), the log-ratios of drop-off/mutation rates (LDRs/LMRs) at each nucleotide position are computed for pairs of control samples to give a null distribution, in order to quantify variability in drop-off or mutation rates observed by chance. LDRs/LMRs are also computed similarly for all possible treatment-control comparisons. Coverage-dependent biases are then removed by applying a variance stabilization transformation. Subsequently, per-nucleotide empirical *P* values are computed for all possible treatment-control comparisons in each condition, by comparing the corresponding log-ratios to the null distribution. DiffBUM-HMM is run on *P* values associated with the two independent conditions as observations, leaving out any nucleotides with missing data. The resulting output is a posterior probability of modification for each nucleotide, ranging from 0 to 1. DiffBUM-HMM reports whether nucleotides were unmodified in both conditions, modified in either of the conditions or modified in both conditions.

### DiffBUM-HMM prediction of structural changes in the 35S pre-rRNA of yeast ribosome synthesis mutants

To test diffBUM-HMM, we first reanalyzed high-throughput structure probing datasets generated from two mutant *Saccharomyces cerevisiae* strains that express structurally distinct pre-ribosomal RNA (pre-rRNA) precursors (33). These ChemModSeq-type (6) high-throughput datasets were selected because (a) the read coverage for the pre-rRNAs analyzed was very high (i.e. *>*10.000 reads per nucleotide) and (b) some of the regions that were predicted to be structurally distinct based on sequencing results have been verified by primer extension (PE) analysis. Since PE analysis is still considered to be one of the most reliable approaches for detecting sites of chemical modification, we used the PE data as ‘ground truth’ for evaluating the goodness of the DRNs predicted by the tools benchmarked in this study, including diffBUM-HMM, deltaSHAPE and dStruct.

In our previous study (34), we validated some of the ChemModSeq results by performing PE analysis on several regions in the 5’ external transcribed spacer (ETS) as well as the 5’ end of 18S (Fig. 2B, regions highlighted in grey; Fig. 3). Here, the ChemModSeq analyses predicted a high concentration of DRNs, which were largely confirmed by the PE data (Fig. 3). DeltaSHAPE and diffBUM-HMM identified many nucleotides as DRNs that also showed differential reactivity in the PE data (Figs. 3A and B). In the 5’ETS, the patterns of the ChemModSeq SHAPE reactivity profiles and the regions verified by PE were very similar (Figs. 3A and B), suggesting that the ChemModSeq high-throughput data for the 5’ETS is of high quality. Both deltaSHAPE and diffBUM-HMM identified DRNs in these regions of the 5’ETS that also appeared differentially modified in the PE data. In contrast, dStruct identified one region in the 5’ETS (nucleotides 462-497; Fig. 3B). One possible explanation for why dStruct reported only one region is because it only searches for DRRs that are longer than a user-specified threshold, whereas diffBUM-HMM calculates posterior probabilities at the nucleotide level. However, reducing dStruct’s search length did not improve the results. A 1-nucleotide dStruct search did not report any DRRs with an FDR of ≤ 0.05. We conclude that, compared to the current gold standard dStruct, diffBUM-HMM detects DRNs with much higher sensitivity and resolution.

**Fig. 2.**
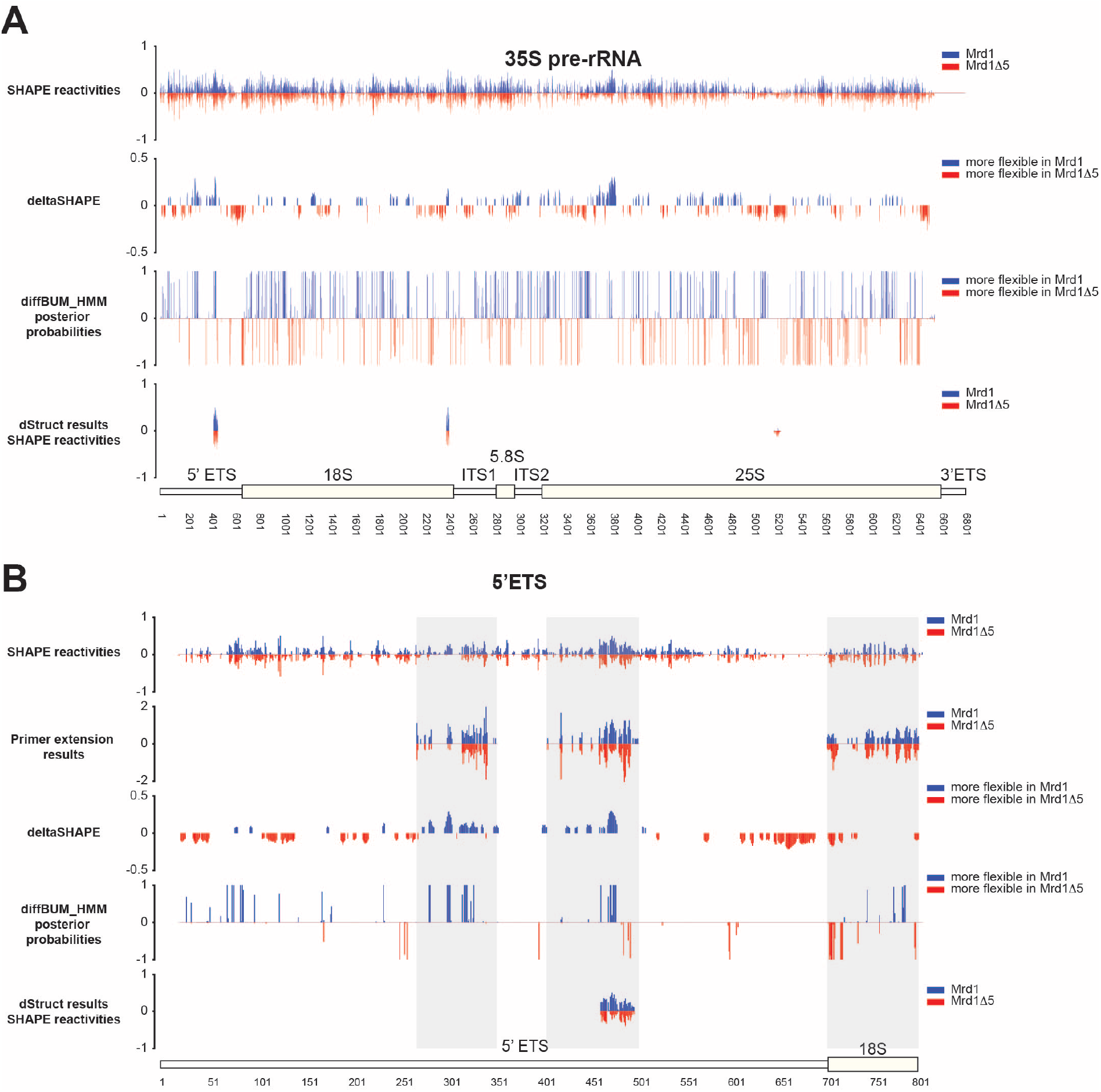
DiffBUM-HMM effectively detects differentially reactive nucleotides in the earliest detectable yeast pre-rRNA precursor. (**A**) The top panel shows the SHAPE reactivities (34) from the first biological replicate of both wild-type Mrd1 and Mrd1Δ5 deletion mutant, which were used to identify DRNs with deltaSHAPE. The deltaSHAPE values were calculated according to (22). For the deltaSHAPE panel, positive values indicate the position of nucleotides that are more reactive in pre-rRNA associated with wild-type Mrd1, whereas negative values indicate the position of nucleotides that are more reactive in pre-rRNA associated with the Mrd1Δ5 mutant. The same data was reanalyzed using the diffBUM-HMM and dStruct algorithms (panels 4 and 5, respectively). For the diffBUM-HMM results, the posterior probabilities for differential states were calculated using the raw counts. For the dStruct analyses, 2-8% normalized RT drop-off rates were used, as recommended by the authors (32). (**B**) Same as in **A** but only for the 5’ ETS and 5’ end of 18S rRNA. We additionally included the results from the PE analysis (panel 2). The grey areas indicate the regions validated by PE analysis.

**Fig. 3.**
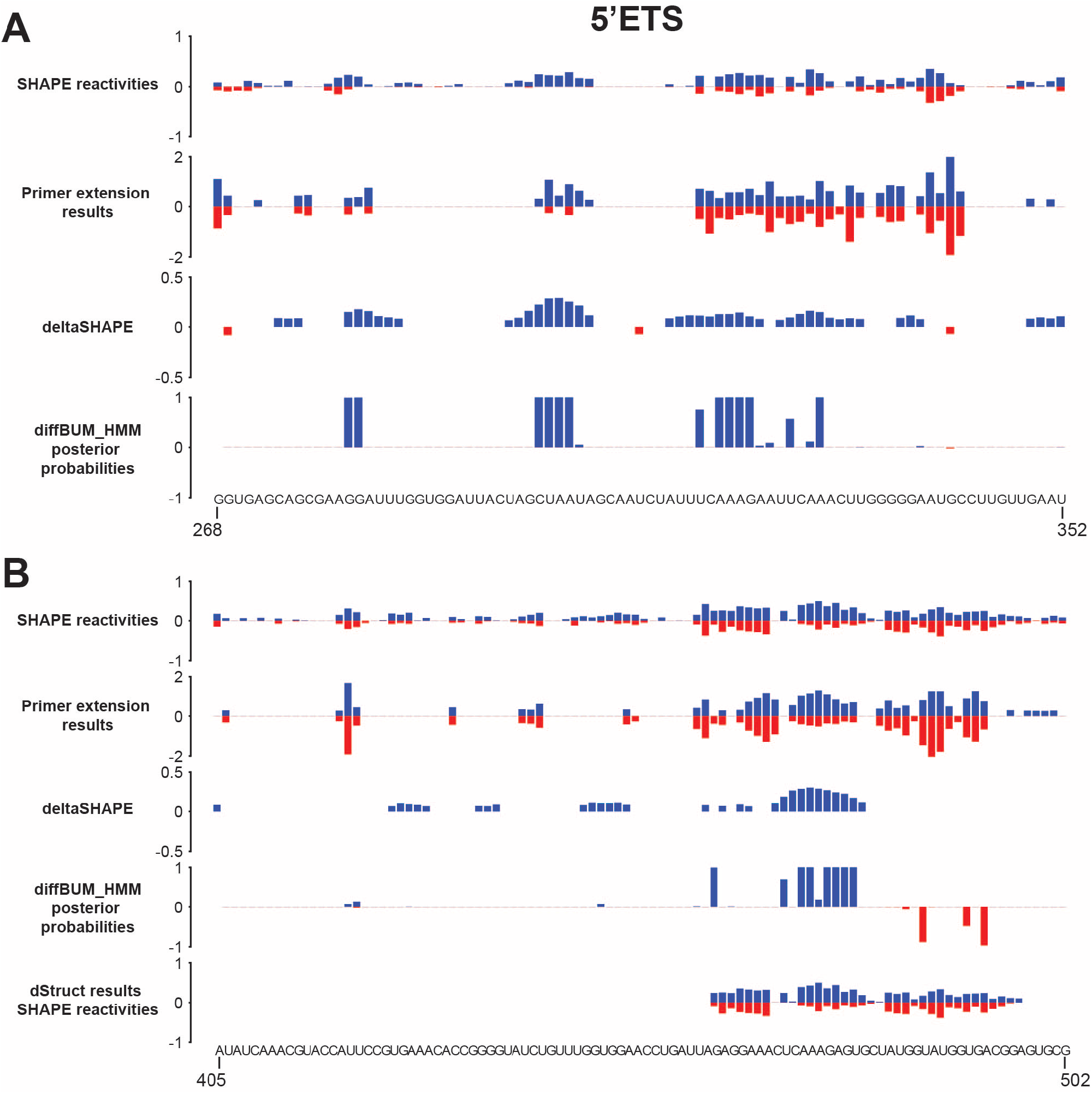
diffBUM-HMM detects differentially reactive nucleotides (DRNs) in the 5’ETS of the 35S pre-rRNA precursor. (**A** and **B**). SHAPE reactivities, deltaSHAPE, diffBUM-HMM and dStruct analysis results for two regions (positions 268-352 and 405-502) within the 5’ETS. The top panel shows the SHAPE reactivities (34) from the first biological replicate, which were used to identify DRNs with deltaSHAPE (deltaSHAPE panel). Positive values indicate the 1M7 nucleotide reactivities in pre-rRNA associated with wild-type Mrd1, whereas negative values indicate the reactivities in pre-rRNA associated with the Mrd1 deletion (Δ5) mutant. The second panel shows the quantification of the PE analysis for these regions. The same data was reanalyzed using the diffBUM-HMM and dStruct algorithms (panels 3 and 4, respectively). For the diffBUM-HMM results, the posterior probabilites for differential reactivity were calculated using the raw counts. For the dStruct analyses, 2-8% normalized RT drop-off rates were used, as recommended by the authors (32). dStruct did not report any DRRs in the region between positions 268-352.

### DiffBUM-HMM calls no false positives in datasets generated from identically treated RNA samples

Despite the fact that deltaSHAPE and diffBUM-HMM were able to detect more experimentally verified DRNs in the 35S dataset, it is plausible that this apparent higher sensitivity is, at least in part, the result of the low specificity of the methods. To test this possibility, we were looking for ways to calculate false positive rates for the diffBUM-HMM algorithm. As the number of nucleotides in the 35S dataset that were verified by PE were too low to perform a meaningful analysis of false positive rates, we reanalyzed published *S. cerevisiae* DMS Structure-Seq datasets generated from *in vivo* experiments that were previously used to assess the false positive rates of all the currently available methods for identifying DRNs (32). These datasets contained biological triplicates of DMS-modified and unmodified mature rRNA samples that were treated identically. Hence, the expectation would be that there would not be any DRNs detected between replicates. We re-analyzed the raw data generated from this experiment and generated drop-off rates for each nucleotide position in the four rRNAs (18S, 25S, 5S and 5.8S). Previously, it was shown that dStruct only called three false positive nucleotides in all the rRNAs analysed here, whereas deltaSHAPE reported a total of 97 false positives (Table 1; (32)). Strikingly, diffBUM-HMM did not report any nucleotide with posterior probability of differential modification higher than 0.4 (Fig. 4A), suggesting that diffBUM-HMM does not call any spurious DRNs in this dataset. This is despite diffBUMM-HMM calling 253 of the 1800 nucleotides modified in all three replicates, which is higher than the value reported for the 18S DMS datasets we previously analysed (134; (27)). DMS preferentially modifies A’s and C’s in flexible and single-stranded regions. Indeed, many of the 18S nucleotides called modified by diffBUM-HMM in all replicates were A’s that were located in single-stranded regions in the 18S secondary structure (Figs. 4B and C). Therefore, we conclude that the data is of good quality and that diffBUM-HMM has a high specificity that is on par with dStruct and RASA.

**Table 1.** Comparison of the capabilities of existing methods designed to detect differential reactive nucleotides in rRNA molecules. The table shows the previously published results of the analyses on the identically DMS-treated yeast rRNA datasets (32) as well as the results from our diffBUM-HMM analysis of these datasets. The column displaying the number of false positives indicates the number of nucleotides that were called differentially modified in DMS chemical probing data.

| Tool | Reference | Inter-replicate variability? | Noise considered? | Detection level? | False positives reported |
| --- | --- | --- | --- | --- | --- |
| classSNitch | (28) | X | ✓ | Regional | N/A |
| PARCEL | (29) | ✓ | X | Regional | 61 |
| RASA | (30) | ✓ | ✓ | Regional | 4 |
| deltaSHAPE | (22) | X | ✓ | Regional | 97 |
| StructDiff | (31) | X | ✓ | Regional | N/A |
| dStruct | (32) | ✓ | ✓ | Regional | 3 |
| diffBUM-HMM | This work | ✓ | ✓ | Regional & Nucleotide | 0 |

**Fig. 4.**
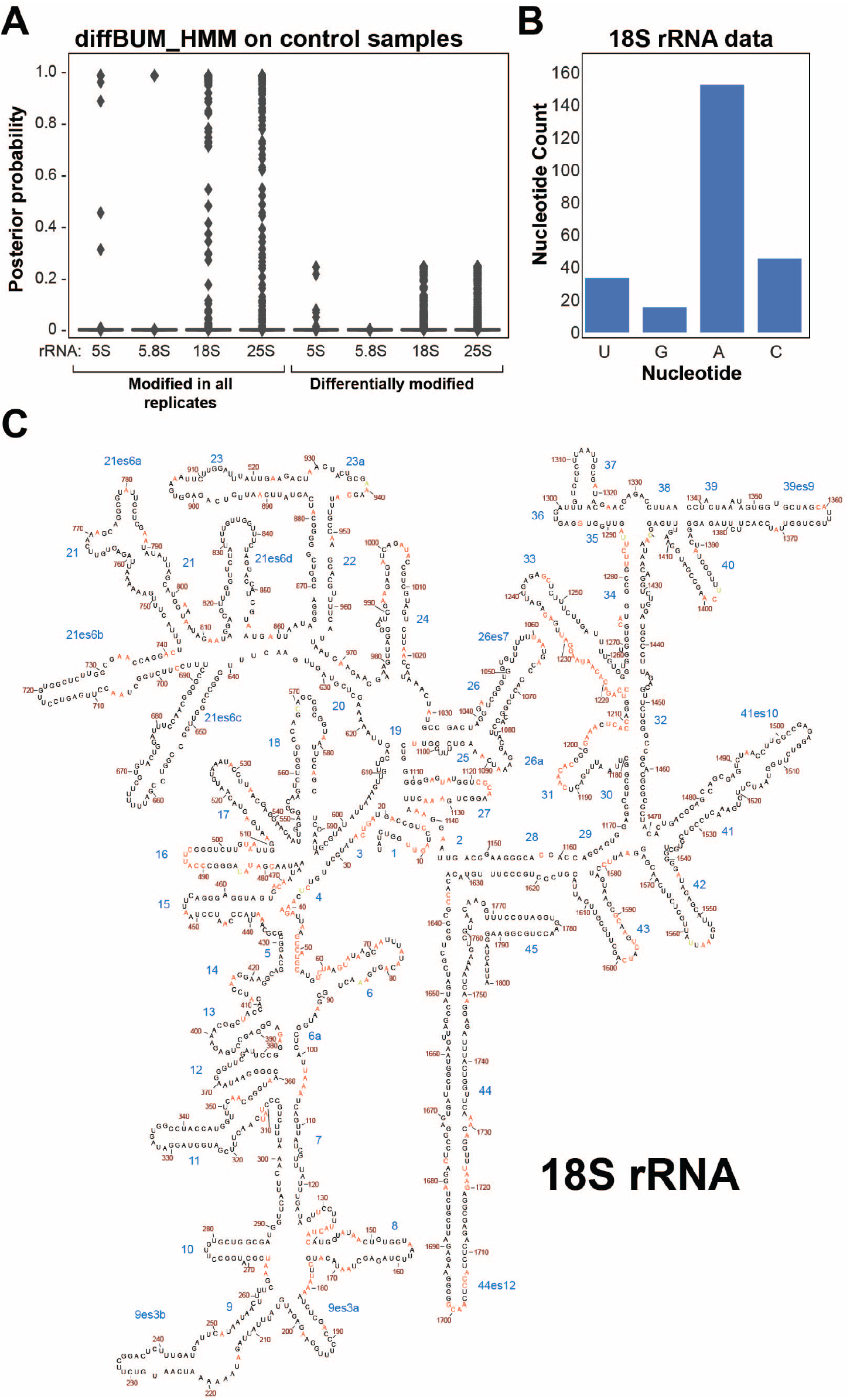
DiffBUM-HMM has a very high specificity. (**A**) DiffBUM-HMM only reports nucleotides with posterior probability of less than 0.4 on identically treated DMS-probed *S. cerevisiae* rRNA StructureSeq datasets. The box plot shows the distribution of the posterior probabilities for each rRNA sample in the control dataset. Shown are the posterior probabilities indicating the likelihood that the nucleotides were called modified in all three replicates (modified in all) or differentially modified between replicates. (**B**) Base composition of nucleotides called modified in all replicates of the yeast 18S rRNA, when considering only nucleotides with posterior probabilities ≥ 0.95. (**C**) Nucleotides that were called modified in all replicates (posterior probabilities ≥ 0.95) are highlighted in red in the secondary structure of the molecule. The names of the helices in the structure are indicated in blue.

### DiffBUM-HMM analysis of differentially probed Xist lncRNA

The earliest studies that reported high-throughput RNA stucture chemical probing analyses relied on the reverse transcriptase falling off the modified RNA once the enzyme encountered a chemically modified nucleotide (6, 8–10, 14, 15). However, by changing the conditions for the RT reaction, one can force a reverse transcriptase to misincorporate non-complementary nucleotides or introduce deletions into the cDNA transcript instead (5). This approach, dubbed SHAPE-MaP (selective 2’-hydroxyl acylation analyzed by primer extension and mutational profiling), maps the site of chemical modification by analyzing the mutation frequency of the nucleotides. To calculate SHAPE reactivities, sequencing data generated from untreated (or solvent treated) RNA and chemically modified denatured RNA are often included. However, it has been suggested that such controls may not be essential for accurately predicting RNA structures (21, 35). Since SHAPE-MaP essentially relies on the counting the number of mutations and diffBUM-HMM relies on count data, we asked whether diffBUM-HMM can accurately detect sites of modification from SHAPE-MaP data. To test this we reanalyzed the mouse Xist SHAPE-MaP datasets (23). The 18-kb Xist lncRNA is essential for X-chromosome inactivation during the development of female eutherian mammals (36). Although previous studies have suggested the importance of RNA structures in specific regions of Xist, the locations and structures of functional domains within Xist are still not well defined. To identify Xist RNA structural features as well as regions occupied by proteins, the Weeks lab recently performed a comprehensive SHAPE-MaP analysis of the Xist RNA that was probed in living cells (*in cell*/*in vivo*) and in protein-free (*ex vivo*) conditions (23). Analyses of these data identified 33 regions in Xist that formed well-defined structures as well as many regions that could be occupied by RNA-binding proteins (RBPs). Importantly, this dataset contained two biological replicates for each condition for SHAPE-treated, untreated and SHAPE-treated denatured RNA samples. As it was unclear whether including the denatured data in our calculation was essential, we performed the diffBUM-HMM with and without normalizing the data to the mutation rates of the denatured RNA samples. An overview of the results is shown in Fig. 5. To compare our data to the deltaSHAPE results, we applied the deltaSHAPE algorithm to the individual replicates (Fig. 5A). When reactivities from the denatured data were not considered, diffBUM-HMM detected 1164 DRNs in the *ex vivo* condition and 188 in the *in vivo* condition (Figs. 5A and C). We generally observed a much bigger difference in the number of DRNs between the two conditions compared to deltaSHAPE (≈ 9-fold with diffBUM-HMM and ≈ 1.4-fold with deltaSHAPE, as shown in Figs. 5A and C). The reason for this is unclear, however, intuitively one would expect that removing proteins from a very large ribonucleoprotein (RNP) complex will substantially increase in the flexibility of the RNA, resulting in more nucleotides being chemically modified on the deproteinized RNA. In this dataset, dStruct was also very conservative with its predictions: with a search length of 11 nt, dStruct reported 8 DRRs with an FDR of ≤ 0.05 (Fig. 5A). Again, shortening the search length to 1 nucleotide did not yield statistically significant DRRs. Remarkably, normalizing the data to the denatured samples further increased the number of DRNs detected by diffBUM-HMM and dStruct (Figs. 5A and C), with dStruct now detecting more DRRs in the 3’ region of Xist (Fig. 5A). This confirms that including data from SHAPE-treated denatured samples can improve the detection of DRNs in SHAPE-MaP data.

**Fig. 5.**
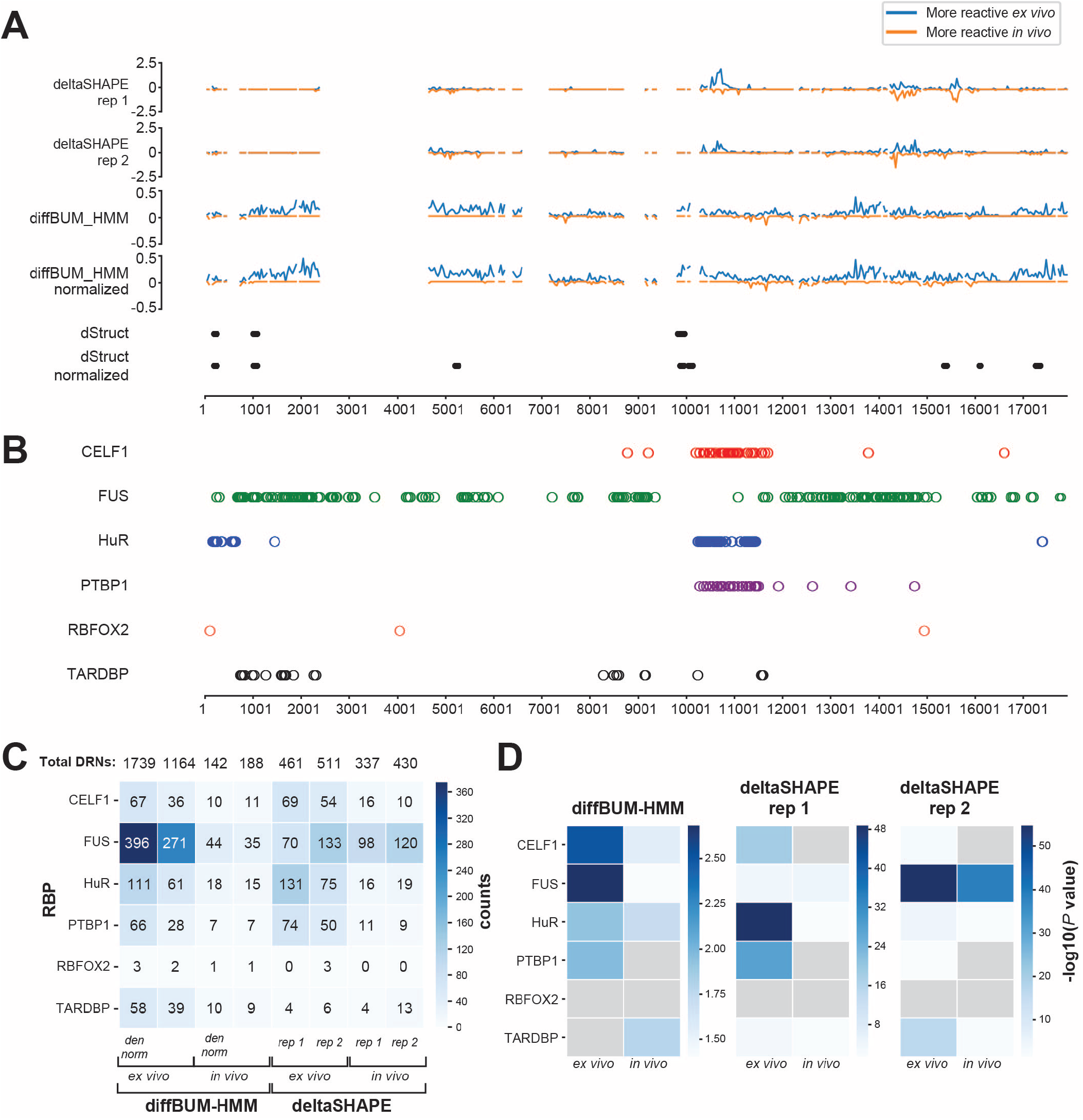
DiffBUM-HMM detects a larger number of differentially modified nucleotides in the *ex vivo* Xist lncRNA data compared to deltaSHAPE and dStruct. (**A**) Shown are the differential reactivities of two deltaSHAPE replicate experiments (23) compared to the diffBUM-HMM and dStruct outputs. The Xist RNA transcript was binned in region of 500 nucleotides and the differential reactivities for each bin is plotted. Regions with negative reactivities are more reactive *in vivo*. Only those nucleotides that according to the deltaSHAPE analyses had sufficient coverage are plotted. The normalized diffBUM-HMM and dStruct panels indicate differential reactivities obtained after normalizing the mutation rates in treated and untreated samples based on the denatured data. (**B**) Overview of RNA-binding sites detected in the Xist transcript, as shown in (23). (**C**) Overview of the number of DRNs that overlap with RNA-binding protein (RBP) binding sites in Xist in the *in vivo* and *ex vivo* data. Total DRNs indicates the total number of DRNs identified by diffBUM-HMM and deltaSHAPE in the datasets. (**D**) Enrichment of DRNs in RBP binding sites in Xist obtained from the CLIPdb database. Statistical significance for enrichment was determined using a hypergeometric test. Colour legend indicates significance level, with binding sites for RBPs that are not statistically significant coloured in grey.

### DRNs detected in Xist using diffBUM-HMM are primarily single-stranded and enriched in protein-binding sites

A key question that we wished to address was whether the large number of additional and unique DRNs detected by diffBUM-HMM in the Xist *ex vivo* data were biologically meaningful. Despite the high specificity of diffBUM-HMM, we could not rule out the possibility that diffBUM-HMM simply called many false positives in this SHAPE-MaP dataset. Deproteinizing an RNP should make sites normally occupied by RBPs more accessible to chemical probes. Therefore, we first asked whether the diffBUM-HMM DRNs were located in protein-binding sites previously identified by crosslinking or RNA immunoprecipitation (CLIP/RIP). The CLIPdb database contains Xist binding sites for a large number of RBPs, including CELF1, PTBP1, HuR, TARDBP, FUS and RBFOX2 (Fig. 5B). Similar to what was previously observed in the Xist deltaSHAPE analysis (23), many of the DRNs in the *ex vivo* data detected by diffBUM-HMM overlapped with RNA-binding sites of these RBPs (Figs. 5B and C). When compared to the deltaSHAPE data, diffBUM-HMM identified m ore D RNs o verlapping w ith F US and TARDBP RNA-binding sites in the *ex vivo* data, whereas the number of *ex vivo* DRNs overlapping with other RBPs was comparable between the two datasets (Fig. 5C). This is presumably because most of the deltaSHAPE signal concentrated around 2-3 regions within the Xist RNA, whereas diffBUM-HMM detected DRNs throughout the transcript (Fig. 5A). We also found that in the *ex vivo* data for both diffBUM-HMM and deltaSHAPE many of the RBP binding sites were statistically significantly enriched for DRNs, with diffBUM-HMM DRNs preferentially enriched in CELF1 and FUS binding sites (Fig. 5D). However, diffBUM-HMM also detected many DRNs outside of these RBP binding sites, which may explain why the *P* values for binding site enrichment are overall lower compared to deltaSHAPE. This is not necessarily surprising since many other proteins bind Xist *in vivo* (37) and therefore diffBUM-HMM could also be picking up binding sites from other proteins in addition to the ones reported in the CLIPdb database. As a second measure for determining whether these unique DRNs could be biologicaly meaningful, we performed a motif search analysis to assess whether enriched sequence motifs could be detected in regions containing DRNs. For this purpose, we grouped together DRNs located within 5nt from each other into genomic intervals, extended these to 30nt and analysed sequence motif enrichment using MEME (38). MEME detected three highly enriched motifs in the CLIPdb binding sites for CELF1, HuR and PTBP1 (Supplementary Fig. S2). Interestingly, similar motifs could also be detected in the diffBUM-HMM and deltaSHAPE data. In the *in vivo* data only a motif resembling the CELF1 binding site was significantly enriched. However, in the *ex vivo* data, sequences resembling HuR and PTBP1 binding sites could be detected. Moreover, diffBUM-HMM again recovered a CELF1-like motif as well as another sequence motif that was not detected in the deltaSHAPE analysis. Thus, these data strongly suggest that the DRNs dectected by diffBUM-HMM are frequently located in or near protein-binding sites.

One possible explanation for why deltaSHAPE calls fewer DRNs in the *ex vivo* data is because it looks within 5 nucleotide windows and only calls a given nucleotide as DRN if at least three nucleotides within that window fit the required criteria. DiffBUM-HMM also assumes that DRNs are present in up to 5 nucleotide stretches, however, the algorithm will call single nucleotide DRNs if it is very clear from the data that only a single nucleotide was differentially modified. Indeed, we found that deltaSHAPE preferentially reports three nucleotide stretches, whereas diffBUM-HMM also frequently reports single nucleotide DRNs (Fig. 6A). If the DRNs uniquely detected by diffBUM-HMM indeed represent real changes in RNA flexibility, one would expect that many of these would be A’s or U’s as these are more frequently located in single-stranded regions such as loops or bulges. This was indeed the case (Fig. 6B). We observed the same trend in the data normalized to the data from the denatured RNA control as well as for those DRNs that were uniquely called by diffBUM-HMM. The deltaSHAPE results were slightly more variable but still showed a modest nucleotide preference. Because the SHAPE reagents used to modify Xist preferentially react with nucleotides in single-stranded or flexible regions, the DRNs called by diffBUM-HMM, including those uniquely detected by the tool, should be primarily located in regions that are predicted to be single-stranded in Xist. Indeed, over 80% of all the DRNs called by diffBUM-HMM in the deproteinized data were located in single-stranded regions (Fig. 6B). A few examples showing DRNs in Xist secondary structures is shown in Fig. 7. In those cases where deltaSHAPE results between replicate samples did not agree, diffBUM-HMM frequently calls the nucleotide unmodified in both conditions. However, as evident from the figures, many of the DRNs reported by diffBUM-HMM were not detected by deltaSHAPE. Collectively, these data suggest that the DRNs detected by diffBUM-HMM in Xist represent *bonafide* changes in nucleotide flexibility that in many cases are located in single-stranded regions and overlap with or are located near protein-binding sites. In conclusion, all the available data strongly suggest that diffBUM-HMM outperforms deltaSHAPE and dStruct in both sensitivity and/or specificity.

**Fig. 6.**
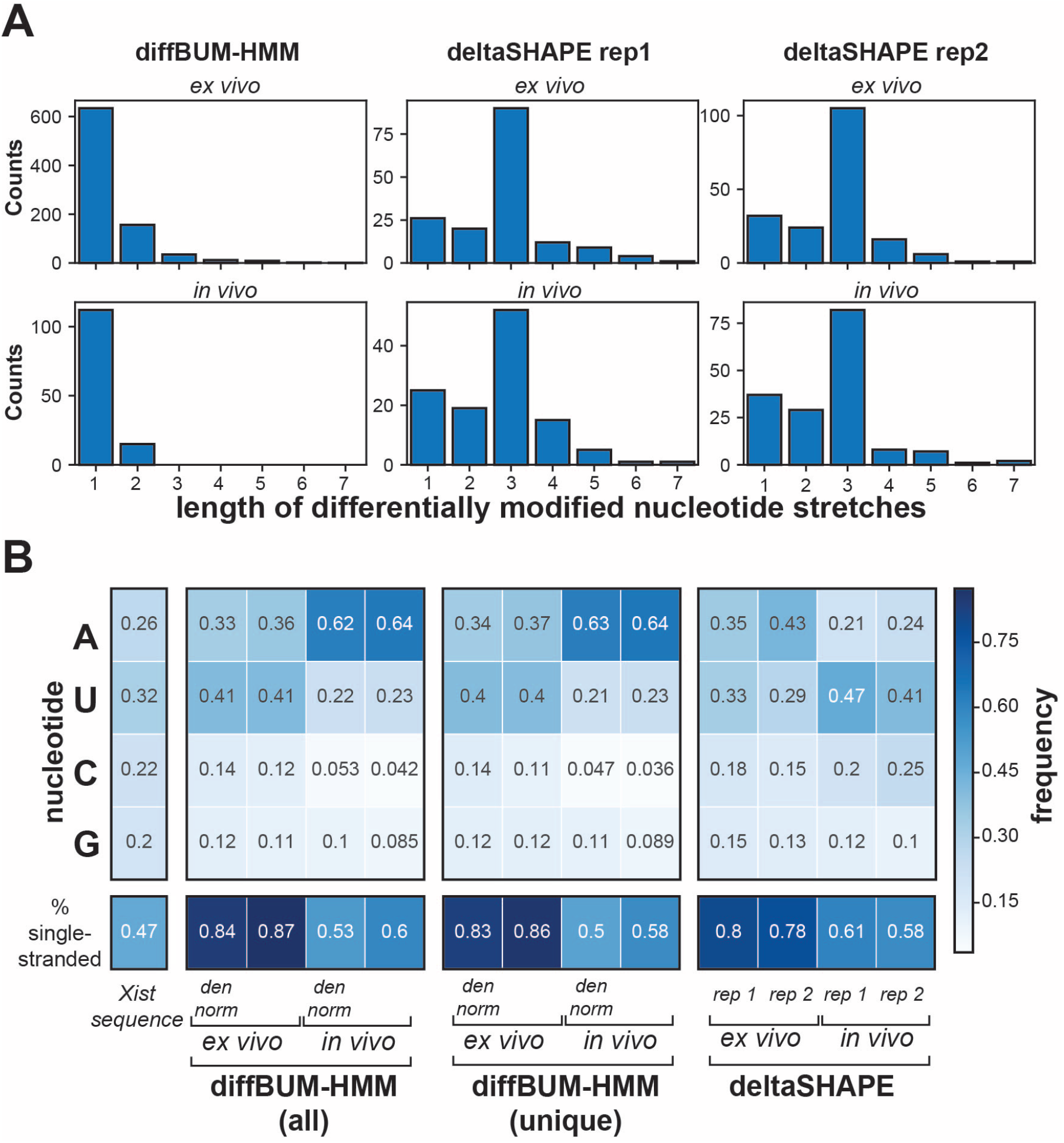
diffBUM-HMM detects more differentially reactive nucleotides (DRNs) in the Xist lncRNA that are preferentially single-stranded A’s and U’s. (**A**) DiffBUM-HMM calls more single nucleotide stretches as DRNs. The barplots show the distribution of the length of stretches of nucleotides that were called DRNs by diffBUM-HMM and deltaSHAPE in the *in vivo* data and *ex vivo* data. (**B**) Shown is the comparison between all the DRNs called by diffBUM-HMM, including the data normalized to the denatured data, those uniquely detected by diffBUM-HMM and the results from the deltaSHAPE analyses on the two replicates individually. DiffBUM-HMM DRNs are mostly A’s and U’s and enriched in regions predicted to be single-stranded in Xist. DRNs identified by diffBUM-HMM are preferentially located in Xist single-stranded regions. ‘Den norm’ indicates the data where we normalized the mutation frequencies of treated and control samples based on the denatured RNA data. ‘DiffBUM-HMM (unique)‘ indicates those DRNs that were uniquely detected by diffBUM-HMM.

**Fig. 7.**
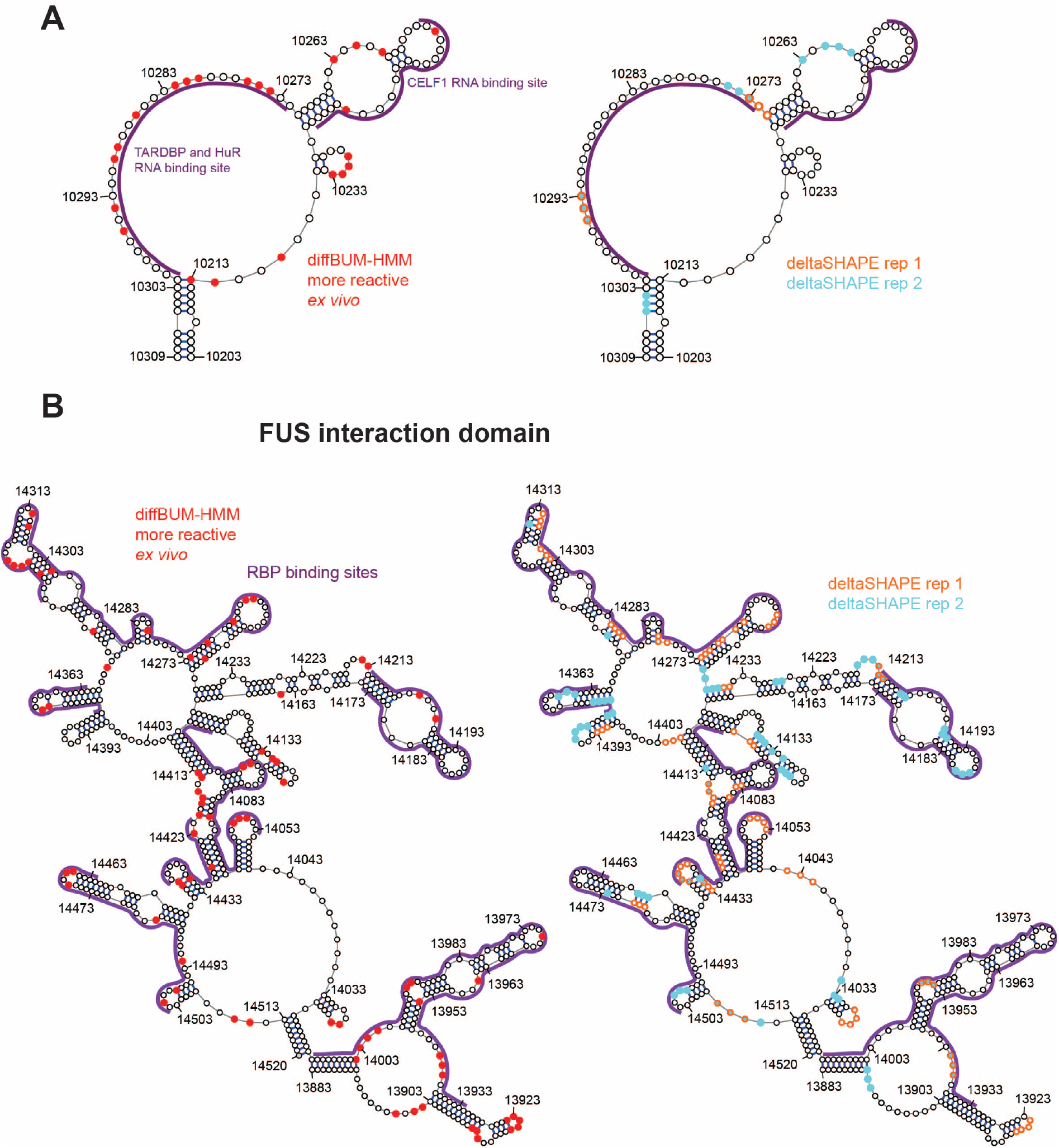
Differentially reactive nucleotides detected by diffBUM-HMM are preferentially localised in single-stranded regions and at the base of stems. (**A**) Shown is a secondary structure for a region in the Xist lncRNA containing CELF1, TARDBP and HuR binding sites. The red dots indicate the nucleotides called modified only in the *ex vivo* data by diffBUM-HMM, while the violet lines indicate binding sites for RNA-binding proteins that were identified by CLIP/RIP. Also shown is the same secondary structure with the deltaSHAPE results from the two replicates individually. (**B**) Same as in **A** but now for the FUS interaction domain of Xist.

## Discussion

Over the past several years there has been an explosion in the number of methodologies that make it possible to analyse RNA structure both *in vivo* and *in vitro*. However, the analysis of the resulting data is notoriously difficult. To be able to extract all the relevant information from the high-throughput sequencing data, many variables need to be taken into consideration. This include sequence coverage, biological variability between experiments (i.e. noise), background signal observed in untreated samples as well as sequence representation bias introduced during the preparation of NGS libraries. Adding to the complexity, research groups have now started focusing on the analysis of RNA structural changes introduced by SNPs or absence of protein binding, etc. This therefore prompted a number of labs to develop bioinformatics tools that would enable users to detect differences in RNA flexibility by comparing datasets generated under different conditions (see Table 1 for examples and references). The Aviran lab recently published a thorough review of the pros and cons of the various methods and tested them on a variety of datasets (32), so we will not discuss this in detail here. However, that study showed that dStruct was the best performing approach, particularly when it comes to specificity. One of the great strengths of dStruct is that it is compatible with a wide variety of RNA structure probing methodologies and takes into consideration biological variability. However, as outlined above dStruct uses a variety of statistical tests to predict DRNs within a certain sequence window. The correction for multiple hypothesis testing that dStruct employs likely also makes the tool conservative with its predictions. Indeed, our analysis of rRNA and mouse Xist SHAPE-MaP data showed that dStruct generally calls few DRRs. This prompted us to develop a tool that was based on a probabilistic graphical model as this should be less vulnerable to problems associated with multiple hypothesis testing. Here we demonstrate that our approach (diffBUM-HMM) is indeed much more sensitive in calling DRNs compared to dStruct on all the datasets tested. However, this high sensitivity does not mean that diffBUM-HMM compromises on specificity: like dStruct, diffBUM-HMM has a very low false positive rate. In fact, our analysis on identically-treated rRNA samples probed with DMS (including rRNAs up to ≈ 3400 nucleotides long; (32)) revealed that diffBUM-HMM did not call any false positives.

Our reanalysis of the Xist data revealed that diffBUM-HMM also called many more DRNs in the *ex vivo* data compared to deltaSHAPE and dStruct, many of which were uniquely detected in the diffBUM-HMM analyses. We believe that the majority of these represent *bona-fide* changes in RNA flexibility: SHAPE reagents preferentially react with single-stranded or flexible nucleotides and over 80% of the DRNs detected in the deproteinized data were in regions that are single-stranded in the Xist secondary structure model (Figs. 6 and 7). Moreover, motif analyses revealed that these DRNs were also enriched in sequence motifs recognised by RNA-binding proteins (Supplementary Figure S2). Our analyses as well as the original Xist SHAPE-MaP paper (23) nicely illustrate how comparing *in vivo* and *ex vivo* conditions can not only help with the detection of differences in RNA structure, but also the identifications of potential protein-binding sites. The observation that specific RNA-binding motifs could be detected in the deltaSHAPE and diffBUM-HMM DRN analyses suggests that it should even be possible to use such data to predict where on the RNA certain sequence-specific RBPs bind.

Although the available evidence suggests that diffBUM-HMM is currently the best performing method for detecting DRNs, it does have a few drawbacks: the method provides a posterior probability for differential modification, which does not inform about how large the difference in chemical reactivity was between the two samples (i.e. no absolute measure of modification). However, to get an impression of reactivity changes, one could plot the average LDRs or LMRs of the different conditions together with the output of diffBUM-HMM. As diffBUM-HMM solely relies on nucleotide count data, it is compatible with a wide variety of high-throughput RNA structure probing methods that either measure RT drop-off or mutations (SHAPE-MaP). However, it is important to point out that diffBUM-HMM will only work well with structure probing libraries that are paired-end sequenced, as in order to quantify and correct for local variability in coverage, the precise start and end position of each cDNA in the library needs to be determined (6, 26, 27). In our analyses we therefore only consider reads that are properly paired (i.e. the forward and the reverse read are mapped within a specified distance on the same chromosome). Hence, diffBUM-HMM will not generate reliable results with RNA structure probing methods that rely on single-end sequencing. Paired-end sequencing is also recommended for SHAPE-MaP analysis as it would enable the selection of high-confidence mutations. DiffBUM-HMM provides a nucleotide-level measure of differential accessibility; however, to obtain insights into global changes in structure, suitably constrained RNA-folding algorithms need to be used. An alternative approach called DREEM was recently proposed (35), which instead relies on *a priori* selecting a set of plausible structures based on the chemical probe reactivity profiles, and then determine relative shifts in abundance of the different structures via a read-clustering approach. Hence, DREEM and diffBUM-HMM perform different tasks but provide complementary information from structure probing data sets; due to this, a direct comparison in performance between the two methods is not straightforward nor necessarily meaningful.

## Conclusion

We describe a novel modelling approach (diffBUM-HMM) for detecting changes in RNA flexibility from high-throughput RNA structure probing datasets. Our results show that diffBUM-HMM exhibits a higher sensitivity compared to the current gold standard dStruct as well as deltaSHAPE and calls very few false positives. We envision that diffBUM-HMM will be very useful for a variety of analytical tasks that pertain to different domains ranging from biomedical science to synthetic biology. DiffBUM-HMM could be used to predict novel RNA regulatory elements from transcriptomic studies, or study the effects of mutations on RNA structure, to pinpoint crucial functional domains in RNA or to identify potential protein-binding sites within RNA. The knowledge from these studies can be used in synthetic biology: to design and screen for regulators that will allow fine-tuning of arbitrary functions in synthetic gene circuits.

## Methods

### Analysis of the ChemModSeq dataset

Drop-off and read counts were generated using the pyCRAC package and the CRAC_pipeline_PE pipeline (https://bitbucket.org/sgrann/kinetic_crac_pipeline). Briefly, Flexbar (version 3.4.0) was used to remove adapter sequences and subsequently the reads were collapsed (py-FastqDuplicateRemover.py) to remove putative PCR duplicates. PyReadCounters from the pyCRAC package was used to calculate drop-off counts and coverage for each nucleotide position in the yeast pre-ribosomal RNAs (pre-rRNAs). These were subsequently fed to diffBUM-HMM.

### DiffBUM-HMM model

Differential BUM-HMM (diffBUM-HMM) is a variant of the beta-uniform mixture hidden Markov model (BUM-HMM) (27), and most of the modelling assumptions made for BUM-HMM also hold for diffBUM-HMM. For example, the transition probabilities are defined based on single- and double-stranded nucleotide stretches derived empirically to be of length 5 and 20, respectively. Emission probabilities follow a beta-uniform mixture model. This design is based on the expectation that nucleotides that are not modified under a given condition are associated with *P* values that follow a uniform distribution (39). On the other hand, accessible nucleotides are associated with *P* values that follow a Beta distribution, as they would exhibit LDR or LMR values that are greater than most values in the null distribution. In practice, adherence to this assumption can be easily monitored by plotting empirical *P*-value distributions as in Supplementary Fig. S3–S5. It should be pointed out that, occasionally, saturation phenomena might result in the presence of two Beta peaks; for example, supplementary Fig. S3 shows a peak of *P*-values near zero, corresponding to nucleotides which have significantly higher drop-off/ mutation rates in treatment (and hence are likely modified), as well as a peak near 1. This additional peak is likely the result of saturation in the treated sample, resulting in abnormally few drop-off reads in unmodified nucleotides; the BUM-HMM likelihood will in any case assign a very low probability of modification to such nucleotides, effectively eliminating any problem that might arise from this mismatch of hypotheses. The *α* and *β* parameters of the Beta distribution were chosen heuristically to be 1 and 10, respectively. This allows to assign approximately equal likelihood under both *P* value distribution hypotheses to nucleotides that have LDR/LMR falling in the highest quantiles of the empirical distribution. The hidden Markov model takes as input continuous regions of nucleotides that satisfy a user-specified coverage threshold (i.e. non-negative threshold for all the experiments in this manuscript) and non-zero LDR/LMR for at least one treatment-control comparison. The novel aspect of diffBUM-HMM is that inference is performed based on two independent observed *P* values, each representing a different condition. The Forward-Backward algorithm is the inference method for computing the posterior marginals of all hidden states.

### Analysis of enriched sequence motifs in regions containing DRNs

To detect enriched RNA binding motifs, DRNs in the *ex vivo* data that were located within a window of 5 nucleotides were grouped in to a single interval, which was subsequently extended to 30 nucleotides using the pyNormalizeIntervalLenghts.py script from the pyCRAC package (40). Subsequently, fasta files containing the Xist sequences associated with these intervals were analysed by MEME (38) using the following bash command: `meme-chip –meme-minw 4 –meme-maxw 10 –meme-nmotifs 20 –meme-p 8 –meme-mod anr –norc –rna –noecho –oc OUTFILE INFILE`.

### Data and Code availability

All the raw and processed data files, diffBUM-HMM R code and Python data processing pipelines used for analysing the data in this study are available from the Granneman Lab GitLab repository (https://git.ecdf.ed.ac.uk/sgrannem) and from Paolo Marangio’s Github page (https://github.com/marangiop/diff_BUM_HMM). PyCRAC is available from PyPI: https://pypi.org/project/pyCRAC/.

## Acknowledgements

We would like to thank Dr. Chantriolnt-Andreas Kapourani and Dr. Alina Selega for their helpful advice with the development of diffBUM-HMM. We would like to thank Prof. Kevin Weeks for providing the Xist SHAPE-MaP mutation count data, read coverage data and CLIPdb coordinates for the RBPs they described in the original paper. We are grateful to Dr. Krishna Choudhary and Prof. Sharon Aviran for their help with the dStruct analysis. We also thank Sergey Belikov for providing the 35S primer-extension data.

## Supplementary Notes and Figures

**Fig. S1.**
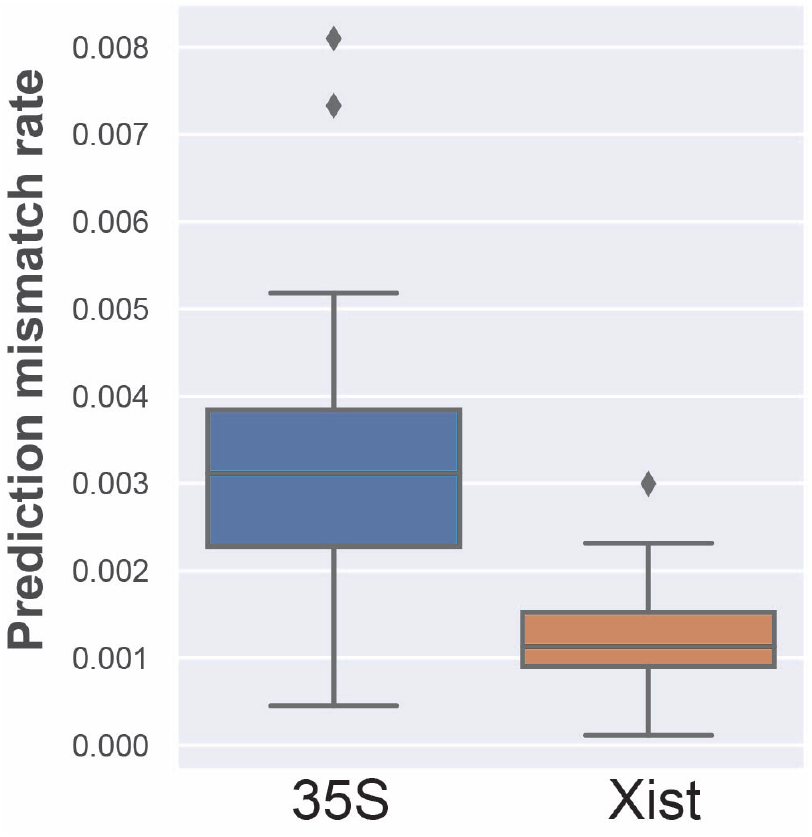
Optimization of diffBUM-HMM transition matrix. Boxplot of prediction mismatch value over 52 transition matrix perturbations for 35S molecule and 61 for Xist molecule. A conservative approach was used in order to adapt the transition probabilities, such that the original values would be respected. Perturbation tests were conducted in order to determine whether the adapted values were indeed optimal. Random Gaussian noise with mean 0 and standard deviation 0.01 was added to the first 3 transition probabilities for each state, while the last transition probability was set such that the 4 values would add up to 1. The resulting posterior probabilities for the differential states (i.e. *hidden state* values 2 and 3) at each position generated when diffBUM-HMM with the noisy transition matrix was applied to the data were then compared against the values outputted with the original, noise-free matrix. A prediction mismatch score can then be computed by dividing the number of incorrect predictions by the number of correct predictions over the entire molecule length, indicating the prediction mismatch associated with an individual noisy configuration of the transition matrix. The prediction mismatch for the 35S pre-rRNA and Xist molecules averaged over more than 50 different, noisy configurations of the transition matrix was smaller than 0.5%. This suggests that the transition matrix configuration used to analyse these datasets was optimal.

**Fig. S2.**
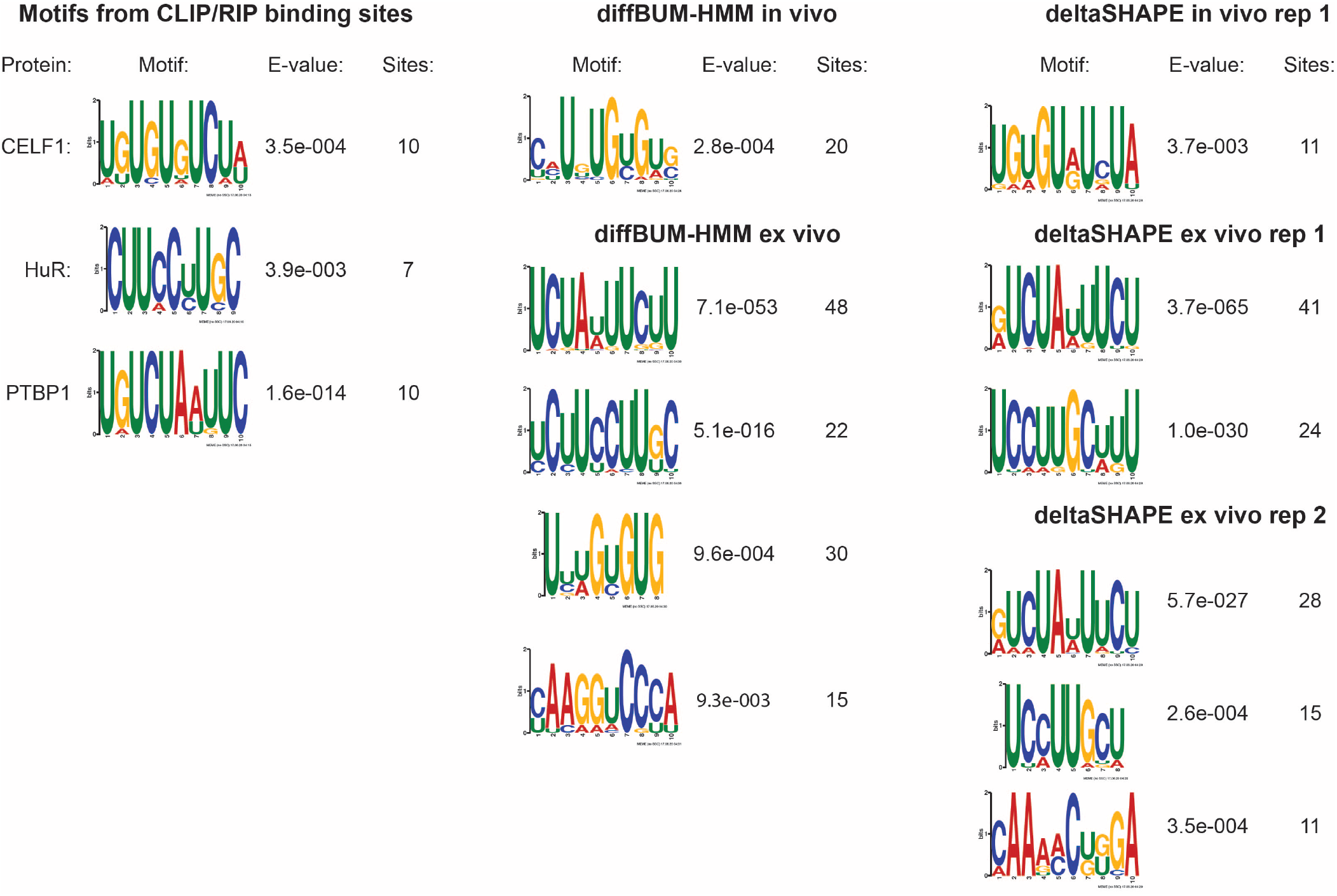
DiffBUM-HMM DRNs are enriched in protein-binding sites. To identify enriched sequence motifs, the MEME tool suite was used (38). The left panel shows the sequence motifs that were detected in the protein-binding sites detected by CLIP/RIP. The middle panel shows the results for diffBUM-HMM and the right panel shows the results for the deltaSHAPE analysis of the individual replicates. Only motifs with an E-value ≤ 0.05 are shown.

**Fig. S3.**
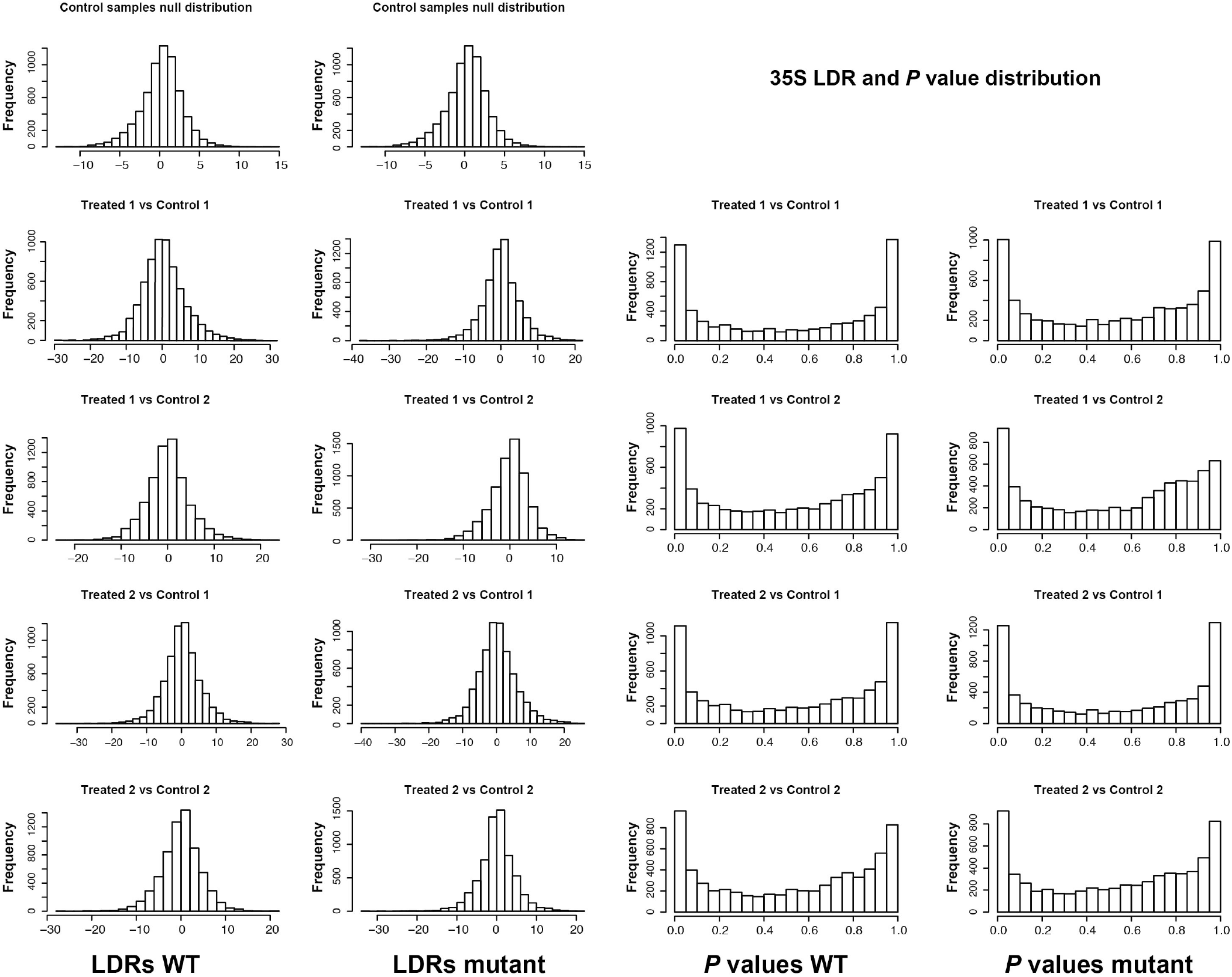
Distribution of log drop-off rate ratios (LDRs) and *P* values for the 35S data. ’Treated’ and ‘Control’ indicate the individual replicates of the SHAPE-modified and DMSO-treated samples, respectively. ‘WT’ indicates 35S pre-rRNA affinity purified using epitope-tagged wild-type Mrd1 as bait, while mutant indicates that the Mrd1 Δ5 variant has been used as bait.

**Fig. S4.**
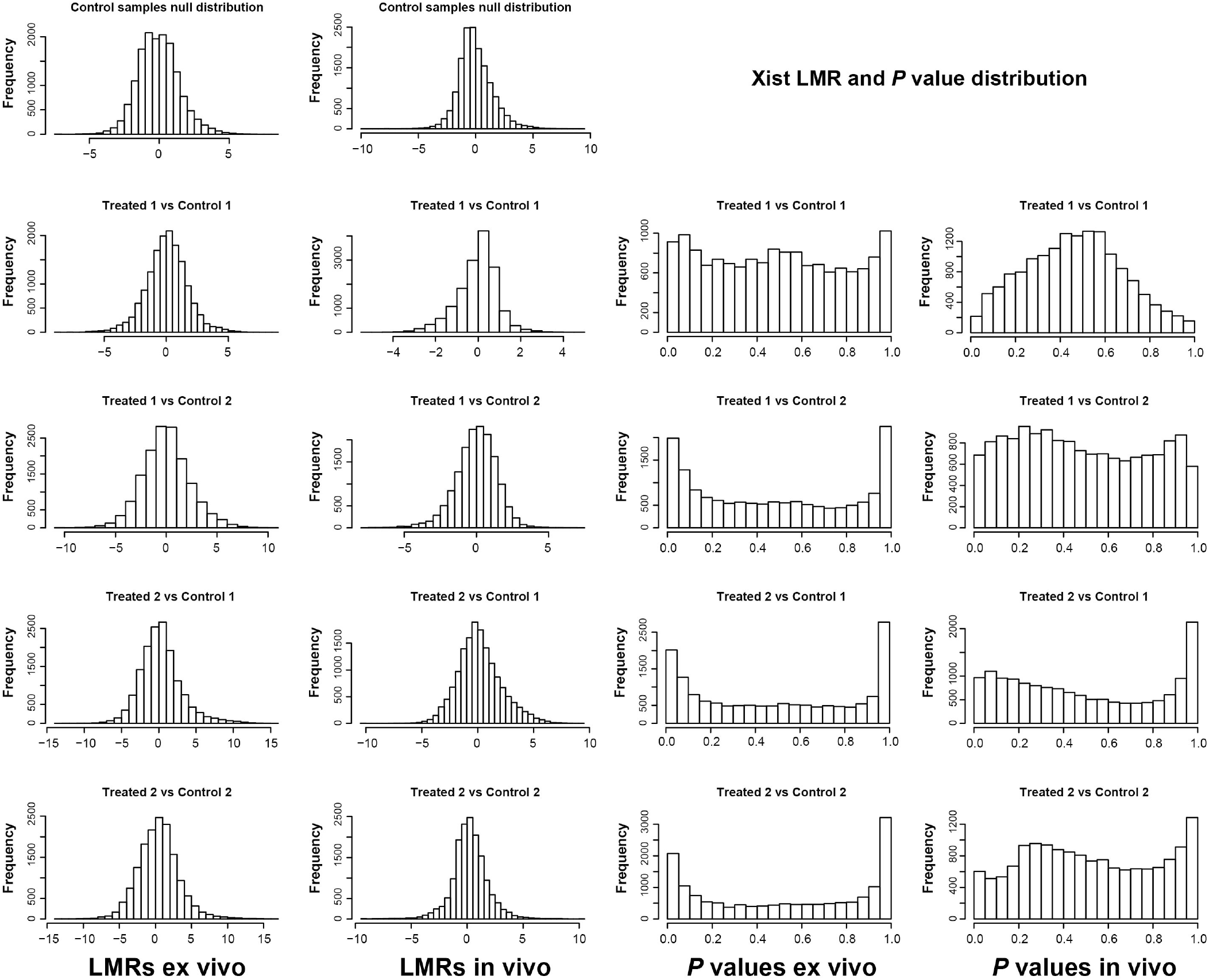
Distribution of log mutation rate ratios (LMRs) and *P* values for the Xist data. ’Treated’ and ‘Control’ indicate the individual replicates of the SHAPE-modified and DMSO-treated samples, respectively.

**Fig. S5.**
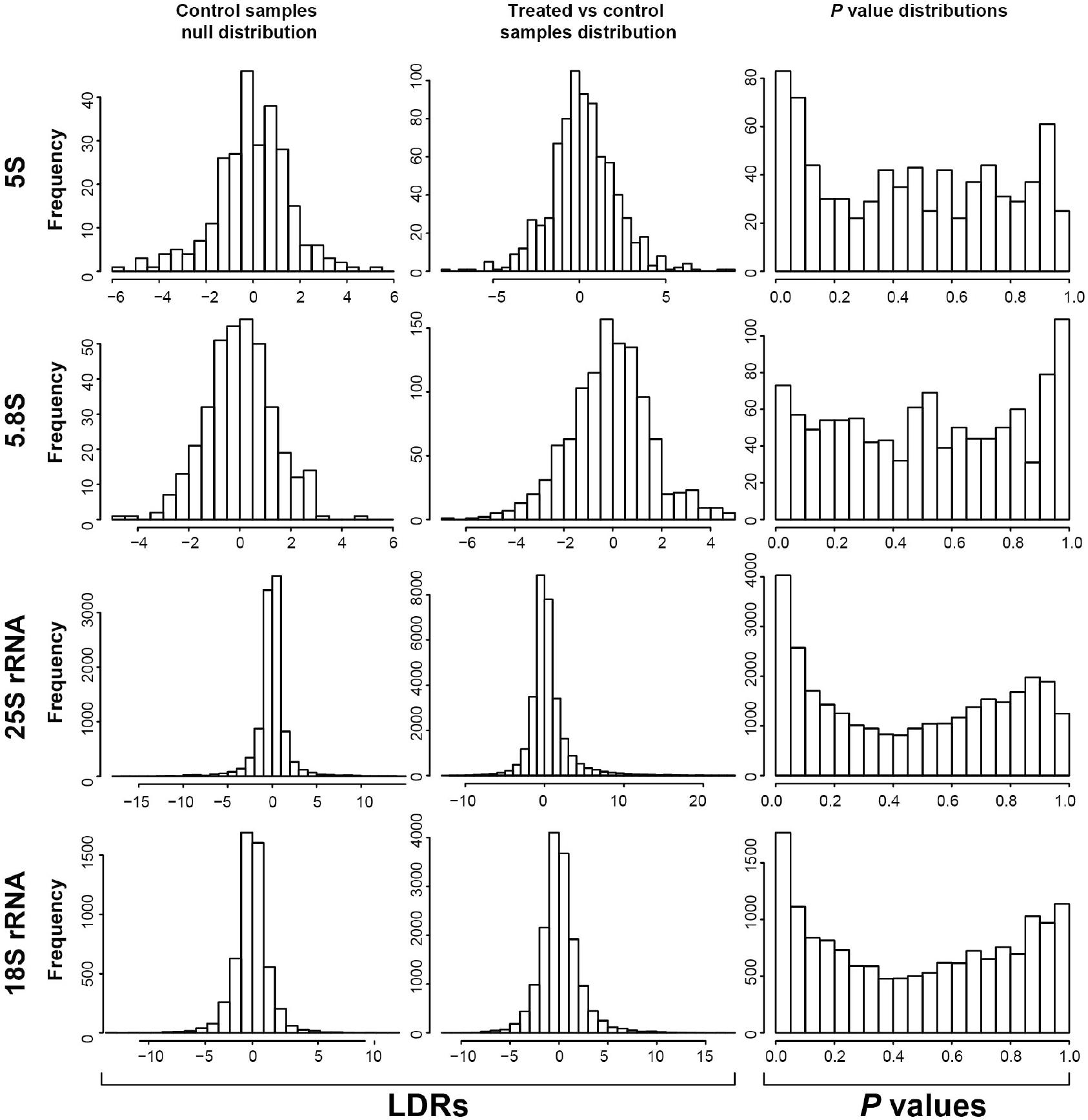
Distribution of log drop-off rate ratios (LDRs) and *P* values for the rRNA control datasets. Since the replicates of the control and treated samples showed a very similar distribution, the data from the different replicates were merged into a single plot. ‘Treated’ and ‘Control’ indicate the samples with and without DMS treatment, respectively.

